# Conditioned place preference reveals tonic pain in Octopus

**DOI:** 10.1101/2020.08.23.263426

**Authors:** Robyn J Crook

## Abstract

Tonic pain is an ongoing, negative affective state arising from tissue damage or inflammation (*1*). Because pain is aversive and its relief is innately rewarding, mammals learn to avoid a context in which pain is experienced, and prefer one where pain relief occurs(*2, 3*). It is generally accepted that vertebrate animals experience pain(*4*), however, there is currently no compelling evidence that pain occurs in any invertebrate(*5*). Here we show that octopuses exhibit tonic pain behavior after subcutaneous injection of dilute acetic acid. In conditioned place preference assays, octopuses avoid contexts in which pain was experienced, prefer a location in which they experienced tonic pain relief, and show no conditioned preference in pain’s absence. Octopuses are thus the first invertebrate shown to experience pain.

**One sentence summary:** A cognitive test demonstrating the emotional component of pain in mammals reveals the first example of pain in any invertebrate.

## Main

Whether invertebrate animals are capable of experiencing pain is the subject of ongoing debate(*6*–*9*). Unlike nociception, which is a simple reflex response, pain is a complex emotional state encompassing distress and suffering, and is generally considered to require a highly complex nervous system(*10*). Discrete pain circuits within the central brain produce two distinct aspects of pain experience; the “discriminative” component, encompassing the location, quality and intensity of pain, and the “affective” component, encompassing the negative emotional state (*11*). Pain is accepted to occur in vertebrate animals, although pain experience that is persistent and ongoing (tonic pain) has to date only been demonstrated in mammals(*1, 12*). Although the evolutionary origins of pain remain unresolved, there is no conclusive evidence indicating the capacity to experience pain has evolved independently in any invertebrate, whose brains are typically smaller and simpler than those of vertebrates(*13*–*15*).

Cephalopod molluscs are extreme outliers in the realm of invertebrate brains; unlike all other invertebrates, their brain size, cognitive ability and behavioral flexibility surpass those of many vertebrates(*16*). Their nervous system is organized fundamentally differently from those of vertebrates, with extensive peripheral control of sensing and movement which seems to occur largely independently of the central brain(*17*). Their large brains and complex behaviors have led to concern for their welfare, and efforts to regulate invasive procedures performed on cephalopods in research laboratories are now established in many nations (*18*). These rules are informed by the untested assumption that cephalopods’ ‘intelligence’ implies they can experience pain, even where no conclusive evidence exists.

Here, a well-established assay for demonstrating the affective component of tonic pain in mammals (*2, 19*) was applied to Octopus. After a single training session in a three-chamber conditioned place preference (CPP) box (Figure 1), octopuses that received subcutaneous injection of dilute (0.5%) acetic acid into one arm (n=8) showed clear avoidance of their initially-preferred chamber, in which they were confined after injection (Fig. 2A, one-sample t-test, p=0.003). Saline-injected animals (n=7) showed no change in their chamber preference before and after training trials (p=0.19). The change in time spent in the initially preferred chamber differed between the two groups (Bonferroni post-hoc test, p=0.006, Figure 2).

**Figure 1.**
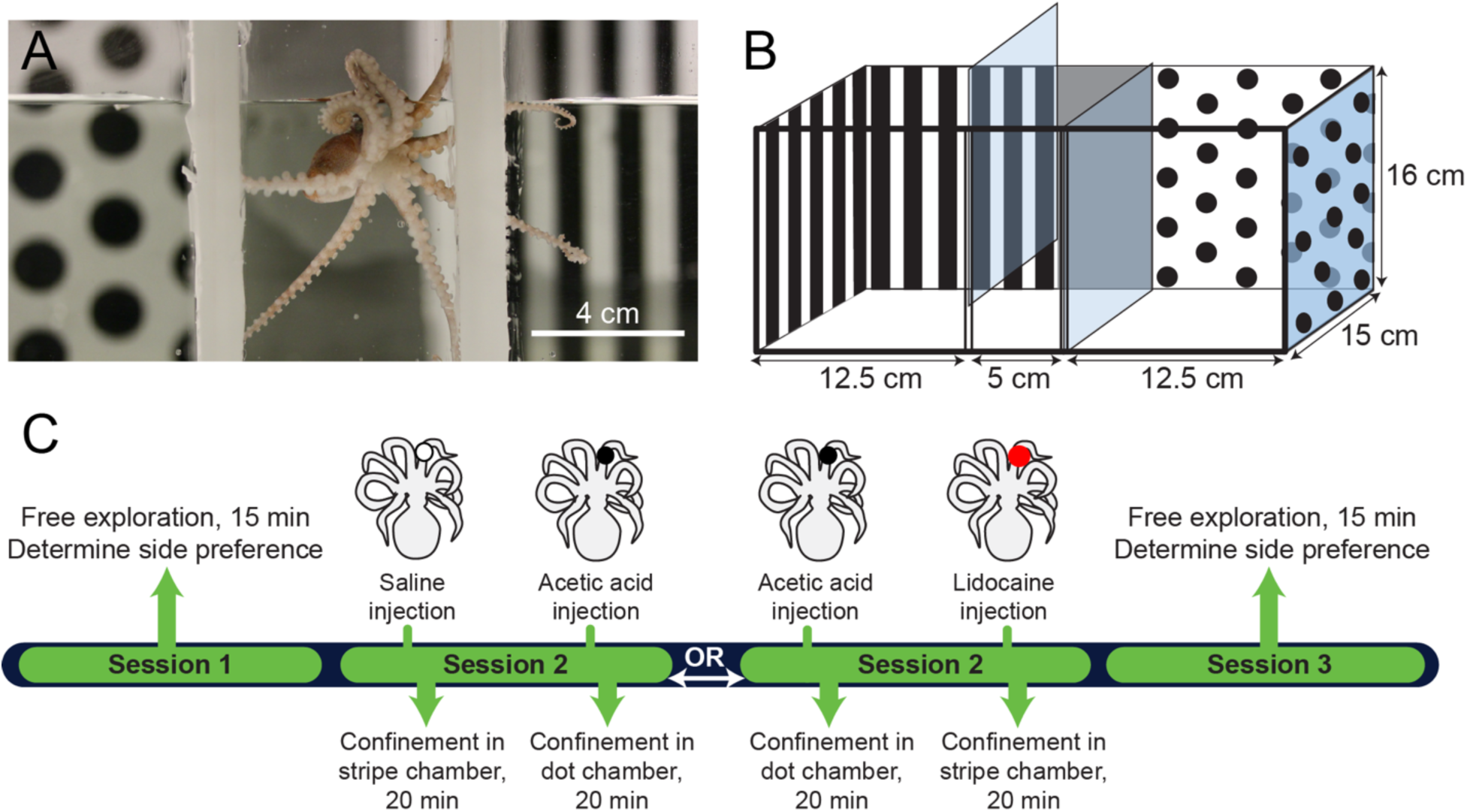
CPP design and timeline. A. *Octopus bocki* in the start chamber of the CPP box. B. Diagram of the apparatus, with pattern shown on the back and sides only for clarity. In experimental trials, visual cues covered all four walls. C. Timeline of an experiment showing sequences for CPA and CPP procedures. In this example, an octopus showed an initial preference in Session 1 for the dot chamber, and is thus trained against initial preference (i.e., the octopus is given AA injection prior to confinement in the dot chamber or lidocaine prior to the stripe chamber). Control sequences (Saline/Saline and Saline/Lidocaine) not shown.

**Figure 2.**
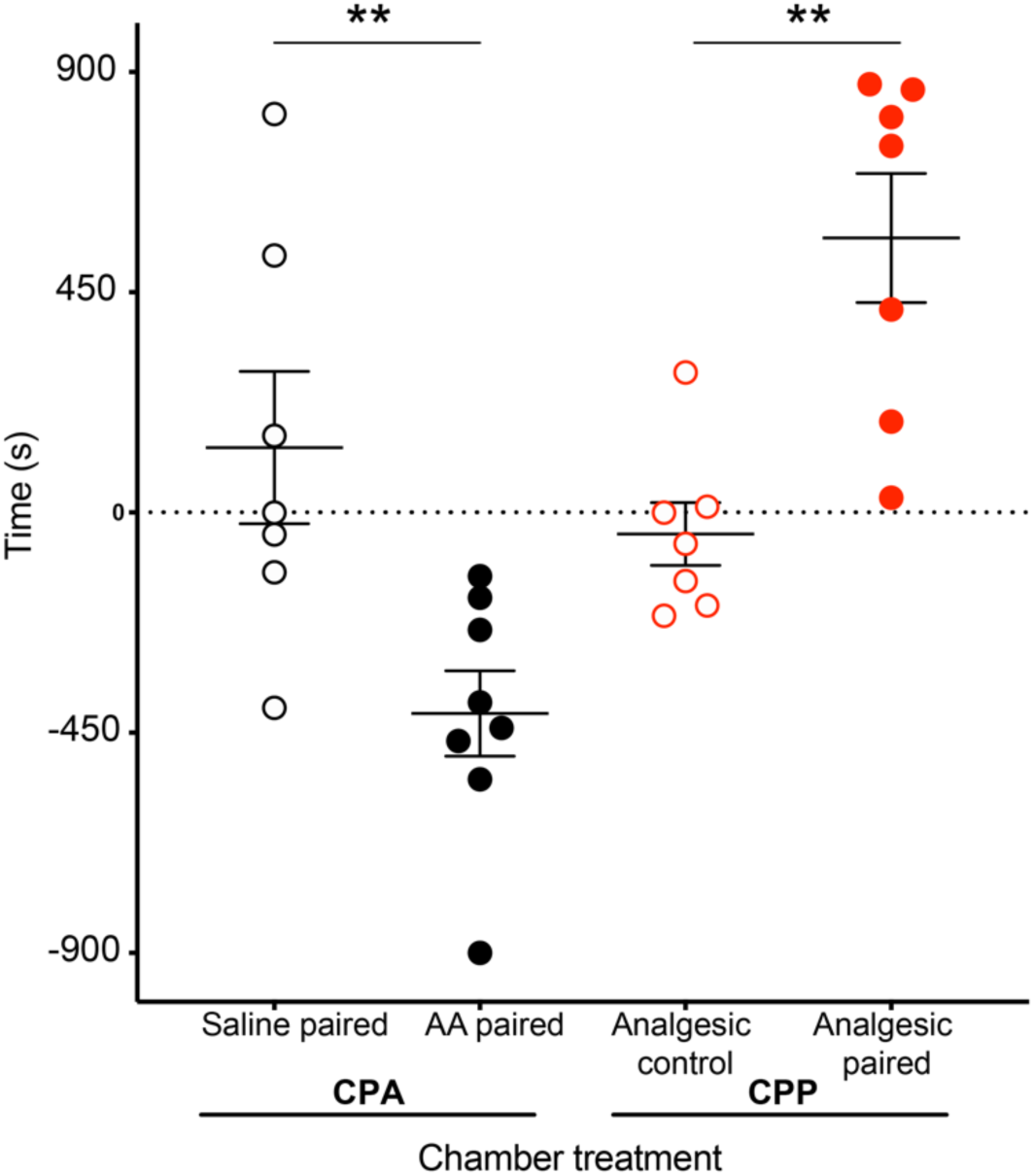
Conditioned Place Avoidance (CPA) and Conditioned Place Preference (CPP) assays reveal the affective component of pain in octopus. In trials where initially preferred chambers were paired with 0.5% acetic acid (AA) injection, octopuses spent less time in their initially preferred chamber in a post-training period of free exploration, compared with octopuses receiving saline. In trials where octopuses received lidocaine over an area of prior injection (either saline or AA), octopuses preferred the chamber paired with lidocaine only if they had previously been given AA injection.

Relief from tonic pain is rewarding, and thus a drug that provides pain relief provides a strong training signal in the presence of tonic pain, but no signal in its absence. Conditioned place preference for a location associated with an analgesic is considered strong evidence for tonic pain in vertebrate animals (*2, 20*). Here, octopuses with AA-induced tonic pain received topical injection of lidocaine (*21*) immediately prior to being confined to the chamber they least preferred in initial preference tests. Lidocaine injection induced strong preference for that chamber in test trials for AA injected animals (Fig 2A, one-sample t-test, p=0.005), but there was no preference for the lidocaine-paired chamber in animals that received saline injection instead of AA (p=0.51), and chamber preference also differed between the two groups (Bonferroni post-hoc test, p=0.003). This demonstrates that lidocaine injection was rewarding to animals only if they were experiencing ongoing pain, and that lidocaine alone is not innately rewarding for octopuses.

While CPP is useful for testing the affective component of pain, it does not necessarily reveal the discriminatory aspect, which includes awareness of the location, quality and intensity of pain (*10, 11*). Point observations of potential pain-associated behaviors (grooming, guarding and concealment) were made at 5-minute intervals during conditioning trials (Session 2), and again 24 h after conditioning trials. All octopuses injected with AA groomed the injection site with the beak for the full 20-minute training trial (Fig 3A), but this behavior was either brief or completely absent in the other groups (Fig. 3B). While wound-directed behavior has been reported previously in octopuses (*22*) and other invertebrates (*23*), the behaviors observed here appear to be specific to acid injection. In all animals receiving AA injection beak grooming resulted in the removal of a small area of skin over the injection site, which was apparent at the conclusion of the 20-minute conditioning trial that followed the injection. This behavior was never observed in animals receiving saline injection or after injection of lidocaine. In other studies of nociception in octopus, arm compression, skin pinch and skin incision induced prolonged beak grooming but never skin stripping (*22*), suggesting that AA injection produced a central representation of pain that was quite different to other injury modalities. Noxious stings, which AA injection likely approximated, are likely encountered by octopuses as they hunt venomous prey (*24, 25*); it is plausible that skin stripping is an injury-induced behavior that has evolved to release injected venom from the skin. This distinct behavior suggests that the octopus central brain is capable of encoding not only the location but also the specifics of pain quality.

**Figure 3.**
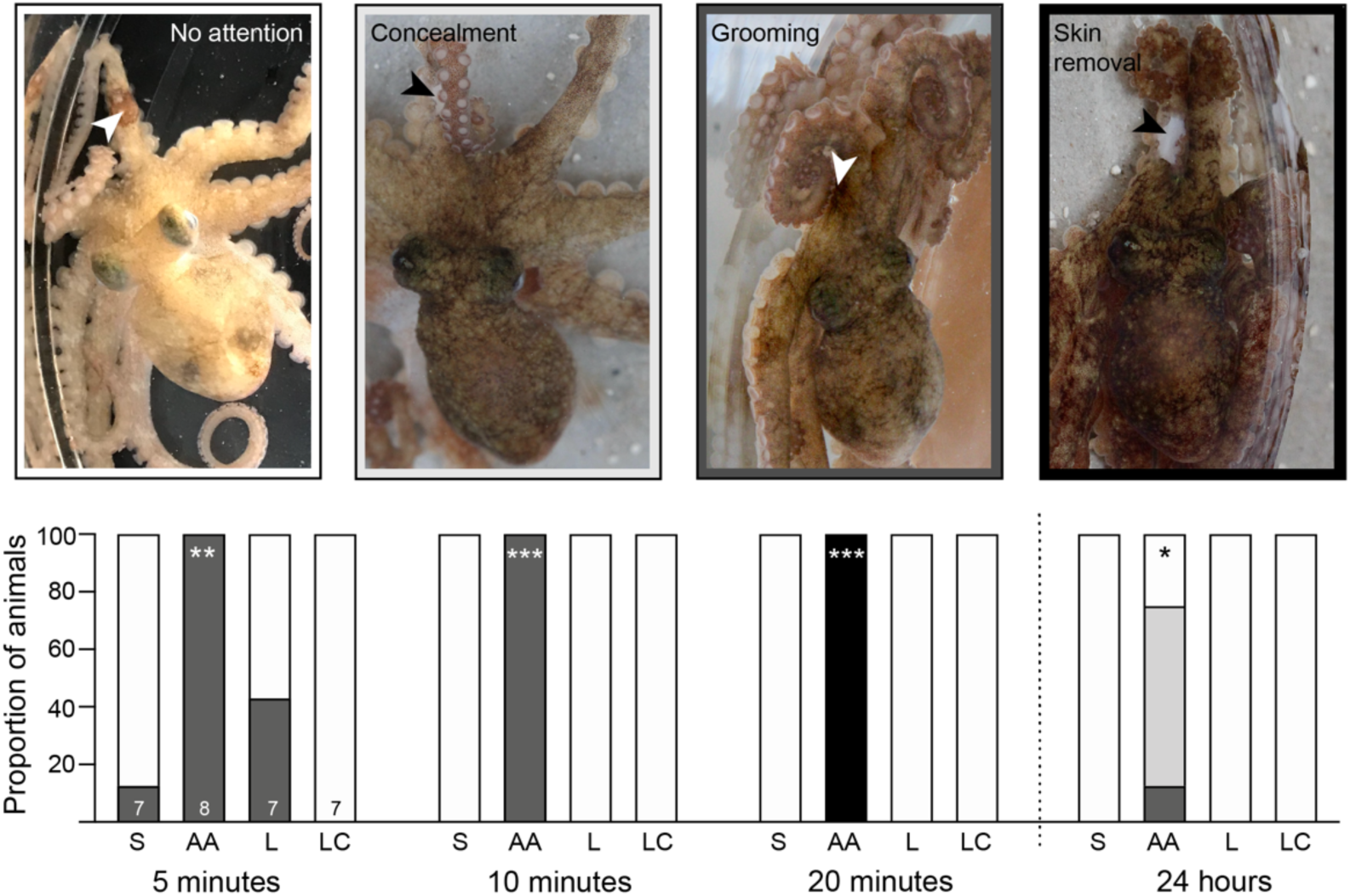
Precise and specific wound-directed grooming behaviors show discriminative pain experience in Octopus. Top panel shows examples of wound-directed behaviors, and colors surrounding each image correspond to shaded frequencies in stacked bars, below. Arrowheads indicate the location of AA injection on the arm. Behaviors were observed during training trials and 24h later. AA-injected octopuses showed sustained wound attention and concealment that persisted for at least 24 hours after AA injection. Skin removal behavior was observed in all AA-injected animals, suggesting a specific representation of acid-induced pain that elicits a highly specific behavioral response. Bar acronyms: AA: acetic acid injection; S: Saline injection; L: Lidocaine injection after earlier AA injection; LC; Lidocaine control (lidocaine injected after earlier saline injection)

Cephalopods are highly unusual in the degree to which higher-order sensory information processing occurs in the peripheral nervous system (*17, 26*). Tonic pain in mammals is driven by sustained activity in primary nociceptors that then drives long-term changes within higher-order, central circuits (*27, 28*). Spontaneous nociceptor firing after tissue injury has also been shown in cephalopods, to date the only invertebrate taxon where this mammalian-like pattern has been recorded (*29*). Whether spontaneous activity in nociceptors drives ongoing excitation of central circuits in the cephalopod brain has not been clear, raising questions of how much the central brain ‘knows’ about noxious sensations in peripheral tissues.

To assess what information the central brain receives about nociceptive stimuli in the arms, electrophysiological recordings were taken from the brachial connectives, which connect the arm nerve cords to the brain and are central to the major arm ganglia situated in the inter-brachial commissure. In a reduced preparation, injection of a bolus of acetic acid subcutaneously into one arm resulted in a prolonged period (>30 minutes) of ongoing activity in numerous recorded units, which was silenced rapidly by injection of 2% lidocaine overlying the site of AA injection (Fig 4). This activity generated within the area of AA infiltration could provide information to the brain about the location of the painful stimulus. Lidocaine injection into the infiltration site also reversed the sensitization of afferent activity evoked by strong mechanical stimulation at and proximal to the injection site, suggesting a role of ongoing afferent activity in promoting evoked pain (hyperalgesia) as well as tonic pain.

**Figure 4.**
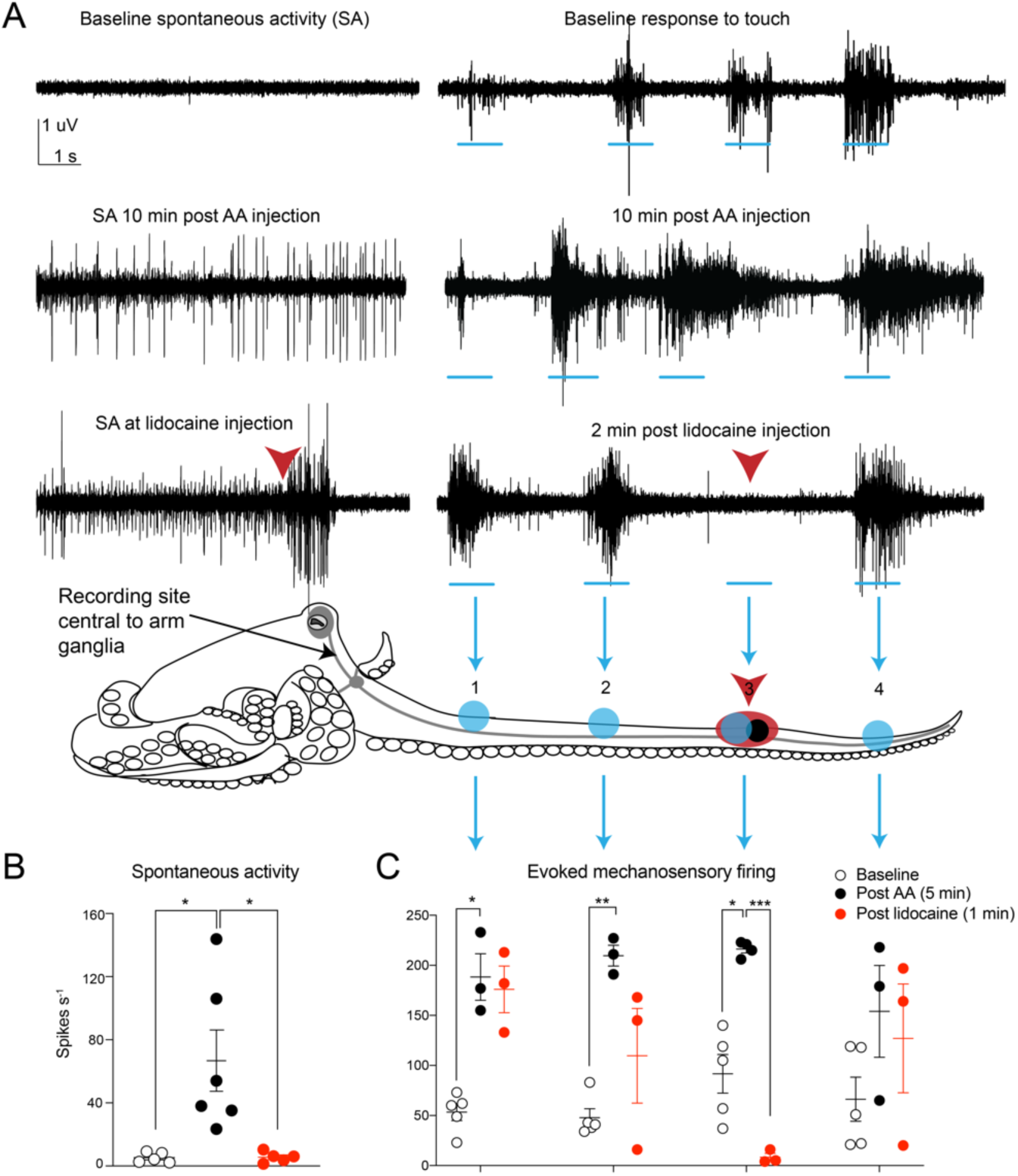
Examples of electrophysiology recordings and summary data showing that nociceptive signal from the arms is available to the Octopus CNS. A. Examples of spontaneous (ongoing) and evoked activity in the brachial connective before and after injection of acetic acid (AA, shown as a black circle at arm stimulation position 3), and at the point where lidocaine is injected locally over the region of prior AA injection (shown as a red overlay of the black circle on the arm at position 3). Note the almost immediate cessation of ongoing activity after lidocaine injection, and the complete suppression of evoked activity in the region where lidocaine was injected at position 3 on the arm of the octopus. B. Ongoing, spontaneous firing in the brachial connective is increased after AA injection and blocked by injection of lidocaine into the same position on the arm. C. Summary data showing responses to touch on the arm at four locations (indicated by shaded blue circles on the octopus body outline). There is clear enhancement of evoked activity after injection that is suppressed by injection of a local anesthetic.

Together, these data reveal the existence of a tonic pain state in octopuses; the first clear example of pain in any invertebrate species and the first example of tonic pain in any non-mammal (*30*). Although a number of previous studies in cephalopods and other invertebrates have shown avoidance learning of a context in which a noxious stimulus was delivered (*31*–*34*), such experiments do not demonstrate the affective component of pain, which relies on higher cognitive and emotional processing. Here, octopuses were able to learn to avoid a visually-specific location that was explicitly unlinked both in time and space from the injection procedure that initiated nociceptor activation, thus the most plausible explanation for the strong place avoidance behavior observed here is that octopuses experiences a state of ongoing (tonic) pain and negative affect after acetic acid injection, which was relieved by local injection of lidocaine into the arm. A common criticism of drug-reversed place aversion is that the chosen analgesic drug is innately rewarding, and its hedonic quality is sufficient to create place preference even in animals in neutral affective states (*19, 35*). The use of lidocaine in this experiment precludes this alternative explanation; lidocaine had no central effect when injected locally, and produced no place preference in octopuses who had not previously received AA injection (and thus were not in pain).

In evolutionary terms, tonic pain is often hypothesized to be adaptive primarily among social species where injured individuals can recruit help from in-group members while ongoing pain reinforces resting and recuperative behaviors (*7*). Additionally, the strong negative affect produced by injury is cited as an adaptive mechanism for reinforcing contextual memory of danger that lasts throughout life. Although the octopus is often described as being “vertebrate-like” in cognitive ability and intelligence, its asocial habits, short lifespan and severe nutritional costs of recuperative inactivity (*36*) argue against the prevailing evolutionary hypotheses cited for pain’s evolution in vertebrates. Instead, the evolution of exceptional neural complexity in cephalopods is typically attributed to their ecological association with complex habitats, niche competition with fish, and their reliance on complex camouflage and signaling behaviors (*37, 38*). How and why pain experience has evolved in cephalopods remains to be understood, and further investigations of the molecular, genetic and anatomical bases of pain in invertebrates will be necessary to shed light on the extraordinary parallel evolution of pain experience in this unique invertebrate clade.

## Supplementary Materials

### Materials and Methods

#### Animals

Adult *Octopus bocki* (Bock’s pygmy octopus, N=29, sex undetermined, average mantle length 14 mm) were obtained from a commercial vendor (Sea Dwelling Creatures, Los Angeles, CA, USA), and housed individually in rectangular tubs (23cm L x 15w x 15.8h, capacity 1900 mL), providing physical, visual and chemical isolation from neighbors. Individual inflow pipes circulated artificial seawater (Instant Ocean, S.G. 1.023, pH 8.1-8.2, 24 Deg C) through each enclosure at a rate of 500 mL/min. Full turnover of water volume occurred every four minutes. Enclosures were located within larger recirculating seawater systems, where water was filtered constantly though physical, biological and charcoal filters. Water quality was monitored daily; ammonia and nitrite were 0 ppm and nitrates ranged up to 20 ppm. Each octopus enclosure contained a bed of crushed coral chips 2 cm deep, three PVC elbow joints of either ½ or 3/4inch, two plastic plants, at least six empty snail and clam shells and two pieces of coral rubble.

Octopuses were fed once per day on a 5mm cube of thawed, frozen, uncooked shrimp (Trader Joe’s brand). Uneaten food was siphoned from the tank once per day during routine tank maintenance. During daily husbandry, octopuses were pre-trained to move from their home tank into a glass beaker to allow tank siphoning, a behavior which also facilitated movement into the conditioning chamber during experiments. Animals were maintained in the laboratory for at least one week prior to being used in experiments, and only animals that were readily accepting food, sheltering normally and were habituated to daily husbandry were used in behavioral experiments.

At the conclusion of behavioral studies, animals that had received painful stimuli were euthanized 24 hours after conclusion of training. The delay was to ensure that the drugs did not induce toxicity or cause death in the acute post-injection period. Octopuses were killed according to established methods (*1*), and tissue was fixed for later use. Control animals were maintained for up to two weeks prior to being used in electrophysiology experiments. Two females in the control group laid eggs within the two weeks and were left to brood their eggs until they died of natural senescence-induced decline.

#### Ethical note

In the United States octopuses are not included in vertebrate animal regulations that govern the use of animals in research. Although no formal approval process occurred, all animal procedures were conducted in accordance with EU Directive 63/2010/EU (*2*), which contains the most stringent requirements for cephalopod research globally. Because the study necessarily involved the use of painful stimuli, sample sizes were calculated to capture moderate and large effect sizes only at a power of 0.8. Post-hoc power analysis indicated 86% power in the CPA experiment and 98% power in the CPP experiment. Procedures, record keeping and reporting were conducted using ARRIVE guidelines (*3*).

### Conditioned place preference (CPP) experiments

#### Apparatus

The CPP arena was made from a modified 9.5 L glass aquarium (Carolina Biological, Item 671226). Two flexible, PVC channels were glued to the sides and bottom of the tank to create holders for two removeable, clear, plexiglass dividers, which when inserted created a three-chamber box (see Fig. 1A&B) with a narrow central start box and two equal-sized end chambers. Visual cues on the tank walls were either black spots (diameter 12 mm, spaced edge-to-edge 6 mm apart) on a white background, or equally spaced black and white, vertical bars (8 mm wide). Walls in the central start box were uniform, 50% grey, and the floor in all three chambers was white. Chamber dividers were clear, but were covered with same-chamber patterns during conditioning confinements in each chamber. The arena was filled with 3L of home tank water, which was not circulated or aerated during trials. Between trials, the water was discarded and tanks were washed inside and out with hot, soapy water to remove any olfactory cues, then rinsed three times with Milli-Q filtered water, sprayed with 70% ethanol solution, and left to dry in bright sunlight. Trials were conducted in an isolated, black-walled room with limited external visual cues. Supplemental, controlled light was provided by a fiber-optic light reflecting diffuse light from the ceiling, which was white. Light level at the water surface was measured with a digital light meter (Dr. Meter LX1010B) at 11 lux. Trials were recorded by a camcorder (Sony FDR-AX33) fitted with a polarized light filter and positioned directly overhead.

#### Drugs

Glacial acetic acid (Sigma-Aldrich, A6283) was diluted in filtered, artificial seawater to produce a final concentration of 0.5% v/v. Sham injections were fASW only. Lidocaine solution (2% HCl) was obtained from A-to-Z Vet Supply (item 515-510212).

#### Procedure

On Day 1, (Session 1, or “Initial Preference Test”) animals were moved from their home tanks and placed into the central start box of the CPP arena. After a two-minute acclimation period, the clear dividers were lifted and octopuses explored freely for 15 minutes. At the conclusion of exploration, octopuses were removed from the CPP chamber in small transfer beakers and returned to their home tanks. Routes taken by each subject were analyzed by Ethovision animal tracking software (Noldus), and end-chamber in which each animal spent the most time (i.e., its initial preference) was recorded. In three cases the octopus did not leave the start box in the first trial. These animals were assigned an initial preference randomly.

The following day, Session 2 (“Training”) comprised two conditioning trials, with the animal confined first in one chamber and then the other. Training was against initial preference, meaning that painful stimuli were experienced in the chamber the animal preferred initially, and neutral or pain-relieving treatments were given prior to confinement in the initially non-preferred chamber.

Prior to the first conditioning trial, animals were removed from their home tank and lightly sedated in 1% EtOH in ASW for handling. Once animals were unresponsive to touch (5-10 minutes after EtOH introduction), one arm was selected for drug treatment. In Experiment 1 (CPA associated with AA injection), 1-2uL of saline was injected about 1/3 along the length of the arm under the dorsal skin, using a 10uL Hamilton syringe and a 30g needle, fitted with a 0.2 micron filter. In Experiment 2, (CPP associated with lidocaine injection), half of the animals received 0.5% AA solution, and half received saline.

Immediately after injection the sedation bath was replaced by running fASW. Animals typically recovered normal behavior within 5-10 minutes. Fifteen minutes after recovery from sedation, octopuses were confined in their initially non-preferred chamber for Experiment 1, and their initially preferred chamber for Experiment 2, for 20 minutes.

At the conclusion of the first 20-minute training trial animals were removed using the standard transfer procedure and allowed to rest undisturbed in small holding tanks for 30 minutes while tanks were washed, dried and refilled with fresh home tank water. After 30 minutes, octopuses were re-sedated for the second injection procedure. In Experiment 1, half of the subjects received 0.5% acetic acid (“AA”) into the arm adjacent to that used for the first injection, while the other half received a second saline injection. In Experiment 2, all the animals received 3uL of 2% lidocaine hydrochloride at the same site as the first injection. Recovery from sedation followed the same procedure as above, and then animals were confined in their initially-preferred chamber for Experiment 1, and their initially non-preferred chamber for Experiment 2, for 20 minutes.

During training, the clear plexiglass divider was replaced with an opaque panel showing the same pattern as the other three chamber walls, thus the pattern in the opposite chamber was completely out of sight during each training. After the second training trial, animals were returned to home tanks. Animal movements were not tracked during single-chamber confinements in the training sessions.

Test trials (Session 3, or “Final Preference Test”) occurred between 5 and 6 hours after the conclusion of the second training trial, on the same day. The procedure was identical to the initial preference test on the preceding day. No drugs or sedation were administered prior to the final training trial.

### Electrophysiology

To ascertain what information the central brain receives about noxious events in the arms, activity was recorded from the brachial connectives, which run between the CNS and the first major ganglion at the top of the arm nerve cord (see Fig. 4). The major arm ganglion lies within the inter-arm commissure, which is a ring linking all the arm that sends signals from one arm to the other. Because there is extensive peripheral processing and sensorimotor integration at the level of the individual brachial ganglia along the arm, and again at the level of the major arm ganglia in the inter-arm commissure, afferent signals recorded from the brachial connectives represent highly pre-processed input into the central brain (*4*). Previous studies have shown that relatively little non-nociceptive mechanosensory information is transmitted centrally from distal arm regions (*5*–*7*), raising the possibility that noxious sensory information is processed entirely in the periphery, without involvement of the central brain.

Octopuses were killed by immersion in isotonic magnesium chloride solution (330mM in Milli-Q filtered water). Ten minutes after respiration stopped, the arm crown was cut from the head and mantle with a scalpel and the brachial connectives exposed by microdissection of overlying tissues. The preparation was pinned tightly into a Sylgard-coated petri dish and the MgCl2 solution was washed off with fASW. One brachial connective was drawn into a suction electrode and the preparation was allowed to rest for 15 minutes. Background firing was recorded for one minute, then a stiff (potentially noxious) von Frey filament (number 5.07, applying 10 g of tip force) was applied to four positions on the arm, moving distally. The stimulation sequence was repeated three times, then the same volume of 0.5% AA used in behavioral experiments was injected into the arm. Background firing and response to the mechanical stimuli were recorded at 1, 5, 10, 30 and 120 minutes in two preparations (data not shown). In six other preparations, 2% lidocaine HCl was injected into the arm at 20 minutes, and background firing and evoked responses ecorded 2 minutes thereafter.

Signals were amplified by an A-M Systems differential extracellular amplifier (model 1700), then digitized and recorded at 10kHZ with a PowerLab 4/35 running LabChart Pro software.

### Data analysis and statistics

CPP: Octopus movements were tracked from recorded video files using Ethovision 13 (Noldus). Examples of tracks and associated data are shown in Fig S1. Time spent per chamber in Session 3 was subtracted from pre-conditioning times spent in Session 1, and all data are expressed as changes from baseline chamber preferences recorded in Session 1. All statistical procedures were conducted in Prism 8.0 (GraphPad). Data distribution was tested with the Kolmogorov Smirnov test and met the assumptions of normality. A single-factor ANOVA followed by planned, post-hoc Bonferroni tests was used to identify between-group differences. To assess whether individual groups’ change in time-per-chamber differed from zero, (a zero value would indicate no change in preference) a one-sample t-test was conducted with an expected value of zero.

Point observations of pain-related behavior were taken every 5 minutes from recorded video footage of training trials. At each point, beak grooming and concealment of the treated area were noted, and frequency per treatment group (proportion of total animals) was compared using Fisher’s exact tests. At the conclusion of training trials, arms were inspected for evidence of skin stripping behavior. Electrophysiology: Spikes above noise threshold were counted using the automated “Spike Histogram” module in LabChart Pro. For each touch, spikes were counted for a 1s period of maximal firing.

Mechanical stimuli were repeated at the same location and timepoint, averaged, and compared at baseline, after AA injection and after lidocaine injection with a repeated-measures ANOVA followed by post-hoc, paired t-tests corrected using the Holm-Bonferroni method (*8*).

All reported p-values are two-tailed. p<0.05 was considered significant.

Data accessibility statement: Raw data associated with each figure are available for download from Open Science Forum under the Project Name “Conditioned Place Preference reveals tonic pain in octopus”.

## Acknowledgements

This study was supported by a CSUPERB New Investigator grant to RJC. Lisa Abbo, DVM provided feedback on the design and conception of the study, and Edgar Walters provided feedback on an initial draft of the manuscript.

## Supplemental Figure

**S1.**
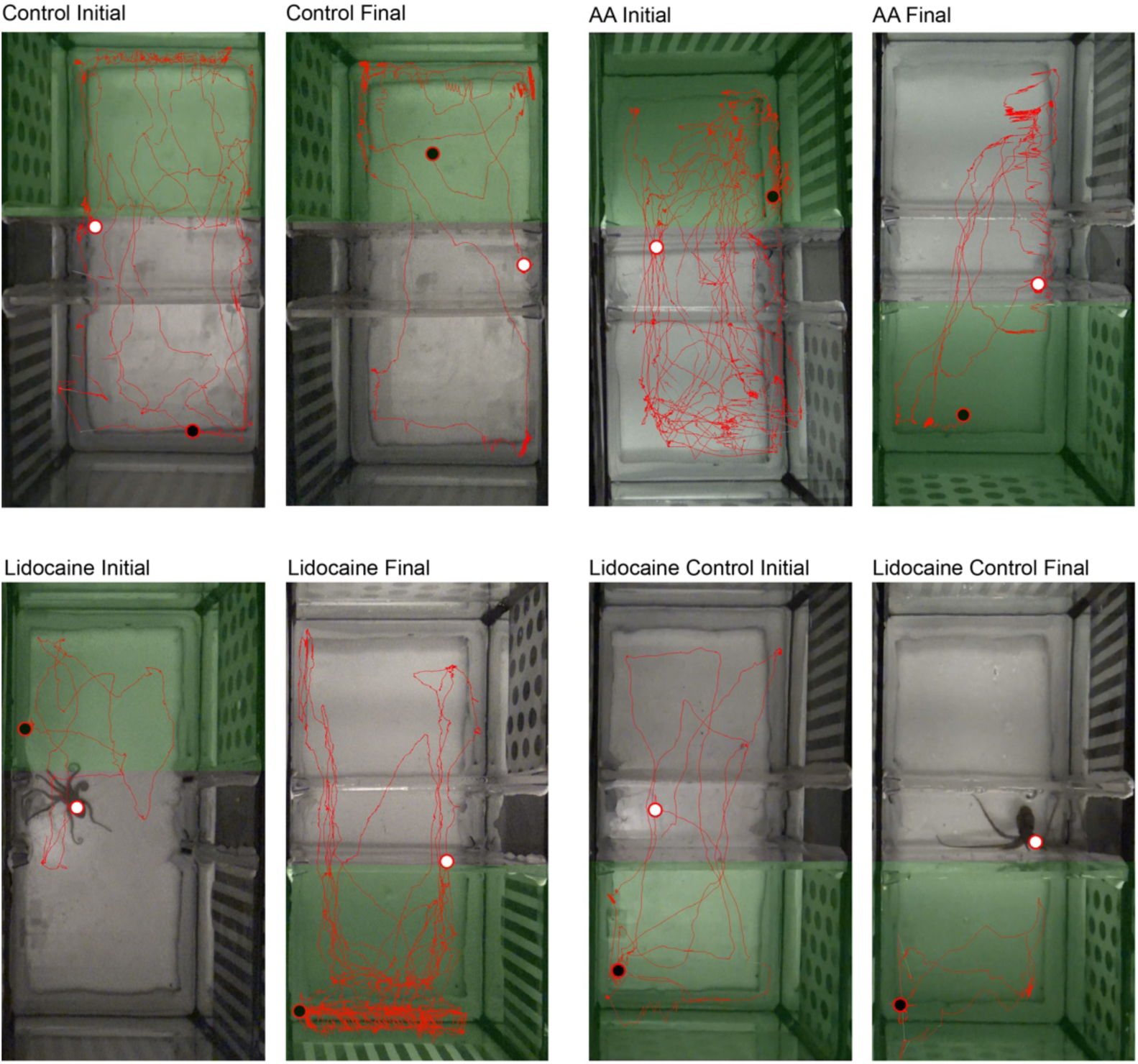
Route maps of representative animals from each treatment group in CPA/CPP assays, generated by Ethovision 13.0 tracking software (Noldus Inc). Routes (red lines) are shown overlaid on a reference image of the chamber for each trial. Start position in the middle chamber is shown by a filled, white circle. Final position is shown with a filled, black circle. The chamber where the octopus spent more time is shaded in green. Octopuses were tracked via center point marker, which was subject to considerable position “jitter” caused by the shift in the computed midpoint as the outline of the animal changed from extended and curled body postures (most notable here in the Control Initial and Lidocaine Final routes). Because this typically occurred along chamber edges it did not affect automatic detection of chamber occupancy.

## Notes

### Competing Interest Statement

The authors have declared no competing interest.

